# Alanine, arginine, and proline but not glutamine are the feed-back regulators in the liver-alpha cell axis in mice

**DOI:** 10.1101/792119

**Authors:** Katrine D. Galsgaard, Sara Lind Jepsen, Sasha A.S. Kjeldsen, Jens Pedersen, Nicolai J. Wewer Albrechtsen, Jens J. Holst

**Author notes:** Shared first-authorship (Katrine D. Galsgaard, Sara Lind Jepsen). Shared last-authorship (Nicolai J. Wewer Albrechtsen, Jens J. Holst). **Correspondence:** Professor Jens J. Holst, Department of Biomedical Sciences and NNF Center of Basal Metabolic Research, Faculty of Health and Medical Sciences, University of Copenhagen, Blegdamsvej 3B, 2200 Copenhagen, Denmark; Or Assistant Professor Nicolai J. Wewer Albrechtsen, Department of Clinical Biochemistry, Rigshospitalet & Novo Nordisk Foundation Center for Protein Research, Faculty of Health and Medical Sciences, University of Copenhagen, Blegdamsvej 9, 2100 Copenhagen, Denmark.

## Abstract

**Aim:** To identify the amino acids that stimulate glucagon secretion in mice and whether the metabolism of these relies on glucagon receptor signaling.

**Methods:** Pancreata of female C57BL/6JRj mice were perfused with 19 individual amino acids (1 mM) and secretion of glucagon was assessed using a specific glucagon radioimmunoassay. Separately, a glucagon receptor antagonist (GRA; 25-2648, 100 mg/kg) or vehicle was administered to female C57BL/6JRj mice three hours prior to an intraperitoneal injection of four different isomolar (in total 7 µmol/g body weight) amino acid mixtures; mixture 1: alanine, arginine, cysteine, and proline; mixture 2: asparatate, glutamate, histidine, and lysine; mixture 3: citrulline, methionine, serine, and threonine; and mixture 4: glutamine, leucine, isoleucine, and valine. Blood glucose, plasma glucagon, amino acid, and insulin concentrations were measured using well characterized methodologies.

**Results:** Alanine (P=0.03), arginine (P<0.001), and proline (P=0.03) but not glutamine (P=0.2) stimulated glucagon secretion from the perfused mouse pancreas. Cysteine had the numerically largest effect on glucagon secretion but did not reach statistical significance (P=0.08). However, when the four isomolar amino acid mixtures were administered there were no significant difference (P>0.5) in plasma concentrations of glucagon across mixture 1-4. Plasma concentrations of total amino acids were higher after administration of GRA when mixture 1 (P=0.004) or mixture 3 (P=0.04) were injected.

**Conclusion:** Our data suggest that alanine, arginine, and proline but not glutamine are involved in the liver-alpha cell axis in mice as they all increased glucagon secretion and their disappearance rate was altered by GRA.

**Graphical abstract:** 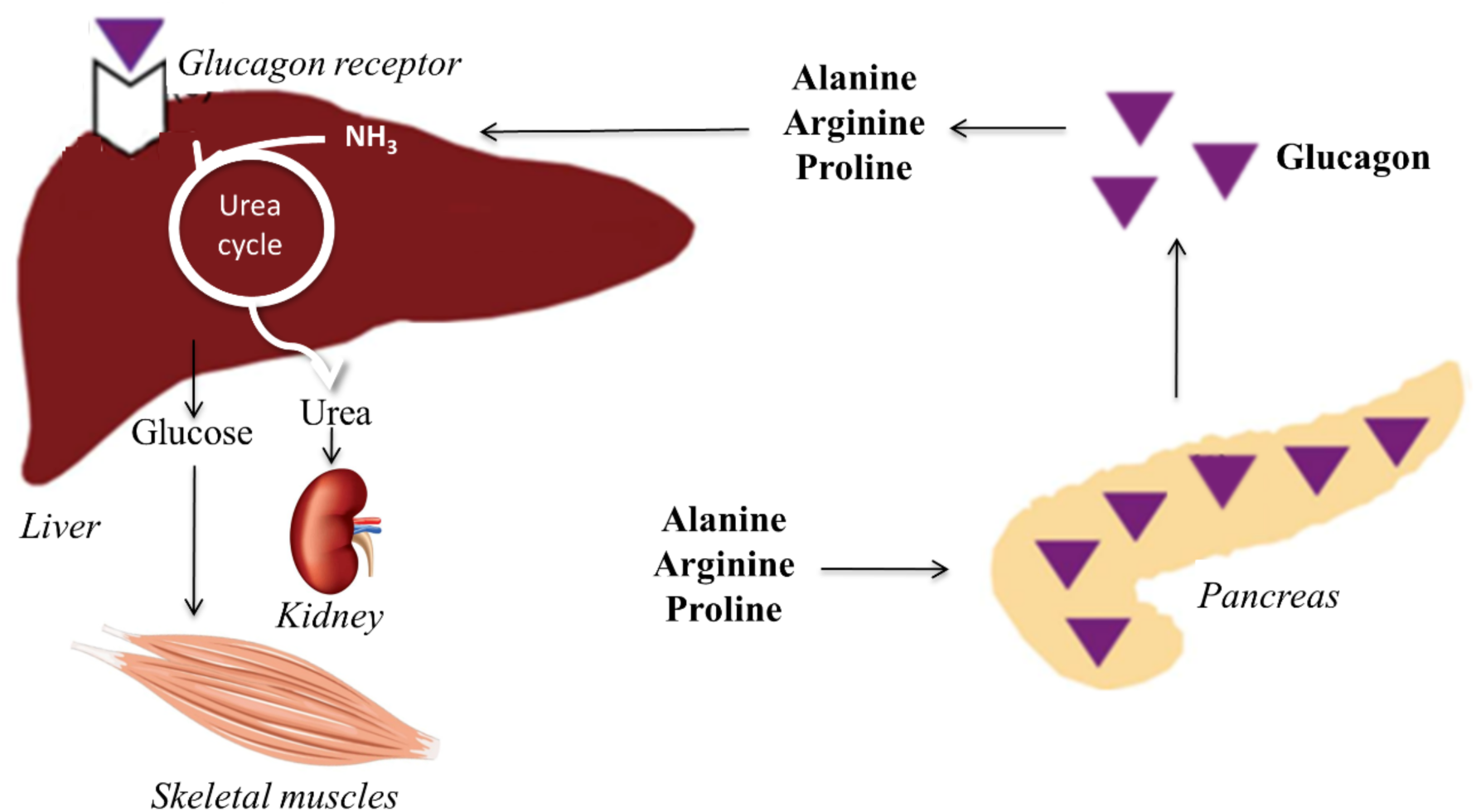

## Introduction

Amino acids stimulate both glucagon and insulin secretion (3, 33, 50, 52, 62). The stimulatory effect of amino acids on glucagon secretion has previously been described to depend on the experimental setting, dose, and type of amino acid administered (3, 17, 52). Arginine is used to assess beta cell function but also increases glucagon secretion (35) whereas the branch-chained amino acids (BCAAs); leucine, isoleucine, and valine, to our knowledge have not been reported to stimulate glucagon secretion (17, 33, 52), with the exception of a single study in which a 20 min valine and leucine stimulation increased glucagon secretion in the perfused rat pancreas (3). By acting on its hepatic seven transmembrane receptor, the glucagon receptor (Gcgr), glucagon increases the hepatic uptake and metabolism of amino acids (5, 22, 24, 25, 28, 45, 47, 49, 56, 63). In line with this, pathologically increased plasma concentrations of glucagon (e.g. glucagon producing tumors) are associated with hypoaminoacidemia (1, 7, 11, 12, 19, 23, 43, 44, 65), which can be extreme, whereas conditions of glucagon deficiency, as observed upon genetic and pharmacological inhibition of glucagon signaling, lead to decreased expression of hepatic amino acid transporters and genes involved in amino acid metabolism, resulting in often dramatically increased plasma concentrations of some but not all amino acids (6, 7, 11, 34, 57). Hyperaminoacidemia may affect alpha cell secretion and proliferation, and thus an endocrine circuit, termed the liver-alpha cell axis, consisting of glucagon and amino acids feedback loops, may exist between the liver and the alpha cells (15, 20, 26, 30, 40, 57, 64). However, which amino acids constitute this feedback loop is not well established, but glutamine has been suggested as a potential candidate (15).

Based on a cross-sectional study in humans (64), we hypothesized that some (e.g. alanine) but not all amino acids are capable of stimulating glucagon secretion directly from pancreatic alpha cells and that their disappearance rate are concomitantly affected by the increased glucagon receptor signaling. To address this, we used a robust experimental setup, the perfused mouse pancreas, to study glucagon secretion in response to individual amino acids, and compared the glucagon responses to the amino acids by injecting four different amino acid mixtures intraperitoneally in conscious mice. Finally, to investigate if the effect of glucagon receptor signaling on amino acid metabolism differs between amino acids, we measured the disappearance of the four amino acid mixtures in the systemic circulation after pharmacological blockade of glucagon receptor signaling.

## Materials and methods

### Animal studies

Animal studies were conducted with permission from the Danish Animal Experiments Inspectorate, Ministry of Environment and Food of Denmark, permit 2018-15-0201-01397, and in accordance with the EU Directive 2010/63/EU and guidelines of Danish legislation governing animal experimentation (1987), and the National Institutes of Health (publication number 85-23). All studies were approved by the local ethical committee. Female C57BL/6JRj mice (∼20 g) were obtained from Janvier Labs, Saint-Berthevin Cedex, France. Mice were housed in groups of eight in individually ventilated cages and followed a light cycle of 12 hours (lights on 6 am to 6 pm) with ad libitum access to standard chow (catalog no. 1319, Altromin Spezialfutter GmbH & Co, Lage, Germany) and water. Mice were allowed a minimum of one week of acclimatization before being included in any experiment.

### Stimulation of the perfused mouse pancreas with individual amino acids

Non-fasted female C57BL/6JRj mice (∼20 g) were anaesthetized by intraperitoneal injection of ketamine/xylazine (0.1 ml/20 g; ketamine 90 mg/kg (Ketaminol Vet.; MSD Animal Health, Madison, NJ, USA); xylazine 10 mg/kg (Rompun Vet.; Bayer Animal Health, Leverkusen, Germany)) and the entire abdominal cavity was exposed. The pancreas was then isolated *in situ* from the circulation as previously described (59). After establishing the perfusion flow through the pancreas, the mouse was euthanized by perforating the diaphragm. The perfusion buffer a modified Krebs-Ringer bicarbonate buffer (pH 7.4, glucose 3.5 mM, and 5% dextran) was gassed with 95% O_2_ and 5% CO_2_. The flow rate was kept constant at 2.0 mL/min, and 1 min effluent samples were collected from the venous cannula using a fraction collector (Frac-920; GE Healthcare, Brøndby, Denmark) and stored at −20°C until analysis. Individual amino acids were dissolved in perfusion buffer reaching a concentration of 20 mM, and were infused intra-arterially at a rate of 0.1 mL/min via a side-arm syringe leading to 1 mM reaching the organ. Each perfusion experiment included four different amino acids. A 5 min arginine infusion was included at the end of the each perfusion experiment and glucagon output in response to this stimulation served as a positive control. The amino acids were obtained from Sigma-Aldrich Denmark A/S, Copenhagen, Denmark (supplementary table 1).

### Pharmacological inhibition of glucagon receptors in mice prior to amino acid stimulation

Non-fasted female C57BL/6JRj mice (∼20 g) received either a glucagon receptor antagonist (GRA; 25–2648 a gift from Novo Nordisk A/S (31)) or vehicle 180 min prior to the experiment. Food was removed at the time of GRA or vehicle administration while free access to water remained. The GRA was dissolved in 5% ethanol, 20% propyleneglycol, 10% 2-hydroxypropyl-β-cyclodextrin (vol./vol.) and phosphate buffer at pH 7.5–8.0 at a concentration of 10 mg/ml, and administered by oral gavage (∼100 µL) as a suspension in a dose of 100 mg/kg body weight as previously described (58). Before GRA administration, blood glucose concentration was measured from tail bleeds using a handheld glucometer (Accu-Chek Mobile, catalog no. 05874149001; Roche Diagnostics, Mannheim, Germany). At time 0 min, referred to as the basal state, blood glucose concentrations were measured and the mice received an intraperitoneal injection of four different amino acid mixtures (table 2) amounting to a total dose of 7 µmol/g body weight, diluted with phosphate buffered saline (PBS) to a total volume of 400 µL. The amino acids were obtained from Sigma-Aldrich Denmark A/S, Copenhagen, Denmark (supplementary table 1). At time 2, 4, 8, and 20 min after the injection, blood glucose concentrations were measured and the mice were subject to a blood collection (∼75 µL) from the retro bulbar plexus using ethylenediaminetetraacetic acid coated capillary tubes (Micro Haematocrit Tubes, Ref. no. 167313 Vitrex Medical A/S, Herlev, Denmark). After the collection of the final blood sample the mice were euthanized by cervical dislocation. The blood samples were centrifuged (7000 rpm, 10 min, 4C°,), and plasma collected and transferred to pre-chilled PCR tubes (Thermowell®, Gold PCR, Corning, NY, USA) and stored at −20C°, until analyzed for glucagon, total L-amino acid, and insulin concentrations. Disappearance rate of the amino acids was evaluated from the incremental areas under the amino acid concentration curve during the 20 min experimental period (_i_AUC_0-20 min_).

**Table 1.** Glucagon secretion from the perfused mouse pancreas in response to individual amino acids (1 mM). The fold change from baseline, graphically shown in Fig. 1, is reported for each amino acid. Significance was assesed by comparing the average glucagon concentration during the amino acid stimulation with the average concentration during the preceding baseline using a paired t-test. Plasma amino acid concentrations prevoiusly reported in (20) for C57BL/6JRj mice are listed and the ratio between the administred amino acid concentration and the plasma concentration is shown. All values, excluding P-values and ratios, are given as mean±SEM.

| Amino acid | Fold change from baseline | Glucagon concentration during stimulation (pmol/L) | Baseline glucagon concentration (pmol/L) | P-value | Amino acid plasma concentrations (μmol/L) | Ratio of given amino acid concentration to average plasma concentration ratio |
| --- | --- | --- | --- | --- | --- | --- |
| Cysteine | 8.4±3.8 | 70±23 | 13±6 | 0.08 |  |  |
| Alanine | 2.1±0.4 | 27±5 | 15±5 | 0.03 | 501±42 | 2.0 |
| Glycine | 1.7±0.3 | 19±4 | 14±6 | 0.2 | 304±14 | 3.3 |
| Proline | 1.7±0.2 | 19±5 | 12±4 | 0.03 | 107±9 | 9.3 |
| Arginine | 1.5±0.1 | 39±6 | 29±5 | <0.0001 | 110±9 | 9.1 |
| Lysine | 1.3±0.1 | 31±10 | 24±7 | 0.2 | 304±17 | 3.3 |
| Phenylalanine | 1.2±0.1 | 42±5 | 37±3 | 0.3 | 80±4 | 12.5 |
| Histidine | 1.2±0.2 | 22±6 | 19±5 | 0.1 | 66±4 | 15.2 |
| Pyruvate | 1.2±0.2 | 28±7 | 32±7 | 0.3 |  |  |
| Serine | 1.2±0.1 | 22±6 | 19±5 | 0.2 | 140±12 | 7.1 |
| Asparatate | 1.2±0.1 | 43±12 | 37±11 | 0.09 | 6±0.5 | 166.7 |
| Asparagine | 1.2±0.1 | 38±6 | 31±2 | 0.3 | 55±6 | 18.2 |
| Glutamate | 1.1±0.1 | 23±5 | 21±6 | 0.3 | 45±3 | 22.2 |
| Threonine | 1.1±0.08 | 27±9 | 24±8 | 0.3 | 173±14 | 5.8 |
| Methionine | 1.0±0.09 | 13±5 | 13±5 | 0.9 | 59±4 | 16.9 |
| Citrulline | 1.0±0.04 | 24±9 | 25±10 | 0.6 |  |  |
| Isoleucine | 1.1±0.1 | 29±6 | 29±6 | 0.7 | 123±10 | 8.1 |
| Valine | 1.0±0.04 | 24±9 | 24±8 | 0.7 | 235±13 | 4.3 |
| Leucine | 1.0±0.05 | 27±7 | 26±6 | 0.7 | 175±12 | 5.7 |
| Glutamine | 1.0±0.05 | 38±14 | 40±15 | 0.2 | 650±19 | 1.5 |

**Table 2.** Amino acid mixtures. The four mixtures were administered *in vivo* as isomolar mixtures, of the four respectively listed amino acids, in a dose amounting to a total of 7 µmol/g body weight.

| Mixture # | Amino Acids |
| --- | --- |
| 1 | Alanine, arginine, cysteine, and proline |
| 2 | Asparatate, glutamate, histidine, and lysine |
| 3 | Citrulline, methionine, serine, and threonine |
| 4 | Glutamine, isoleucine, leucine, and valine |

### Biochemical analysis

Plasma concentrations of glucagon were measured using a validated (66) two-site enzyme immunoassay (catalog no.10-1281-01; Mercodia, Upsala, Sweden). Plasma concentrations of insulin were quantified using an enzyme-linked immunosorbent assay (catalog no. 10-1247-10; Mercodia AB, Uppsala, Sweden, respectively). Plasma concentrations of total L-amino acids were quantified using an enzymatic assay (catalog no. ab65347 Abcam, Cambridge, UK). Glucagon concentrations from the perfused mouse pancreas were measured using a C-terminally directed assay, employing antiserum 4305, which only detects glucagon of pancreatic origin (48) in all efluent samples (1 min), collected from the venous cannula allowing detailed and reliable determination of secretory dynamics.

### Statistics

For analysis of the perfusion data, the average glucagon output over the 10 min periods of amino acid stimulations were compared with the preceding 5 min ‘wash periods’ (this five min period is reffered to as baseline), and the fold change from baseline was calculated by dividing the avarage glucagon concentration during the amino acid stimulation with the average glucagon concentration during baseline. Significance was assesed by comparing the average glucagon concentration during the amino acid stimulation with the average concentration during the preceding baseline using a paired t-test. For analysis of the *in vivo* data, the incremental area under the curve (_i_AUC_0-20 min_) was calculated using the trapezoidal rule after adjusting for baseline values. When two groups were compared significance was assessed by unpaired t-test. When more than two groups were compared significance was assessed by one-way ANOVA followed by a post hoc analysis to correct for multiple testing using the Sidak-Holm algorithm. In all tests, P<0.05 was considered significant. Data are presented as mean±SEM unless otherwise stated.

## Results

### Alanine, arginine, and proline but not glutamine stimulate secretion of glucagon from the perfused mouse pancreas

To assess the effect on glucagon secretion of the individual amino acids independent of the liver, we used the isolated perfused mouse pancreas. Nineteen amino acids (tyrosine and tryphtophan could not be dissolved in the perfusion buffer at a sufficiently high concentration and were excluded) and pyruvate (suplementary table 1) were each administered for a period of 10 min followed by a 15 min “wash period”. Each experiment included four different amino acids, and to control for inter-experiment variation and to confirm that the organ was still responsive we included an arginine stimulation at the end of each experiment. Two examples of the pancreas perfusion experiments are shown in Fig. 1, all perfusion results can be found in suplementary figure 1.

**Fig. 1.**
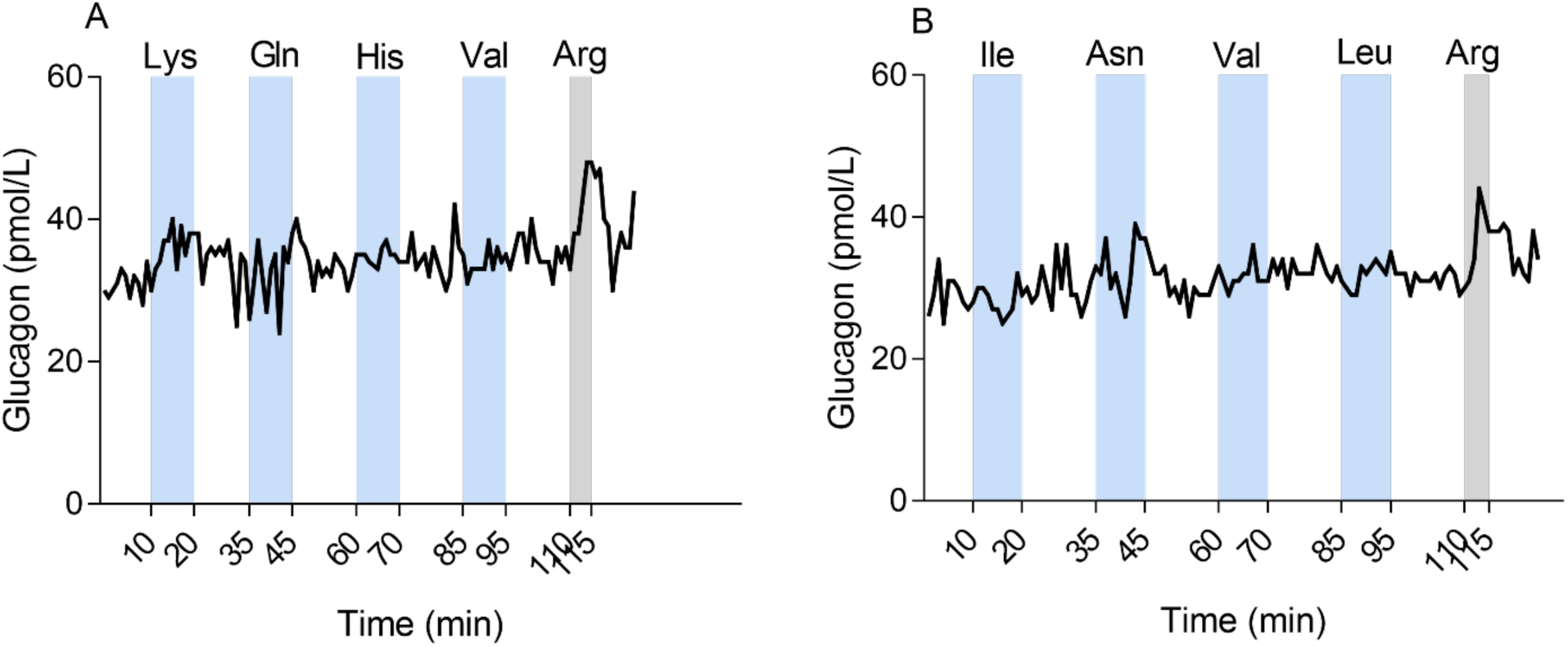
Pancreas perfusion experiments. Glucagon concentrations in effluent from the perfused pancreas of non-fasted female C57BL/6JRj mice in response to; (*A*) lysine (Lys), glutamine (Gln), histidine (His), valine (Val), and arginine (Arg) and; (*B*) isoleucine (Ile), asparagine (Asn), valine (Val), leucine (Leu), and Arg. The amino acids were administered by intra-arterial infusion at a concentration of 1 mM and the glucose concentration was kept at 3.5 mM. The blue bars indicate the 10 min amino acid stimulation and the grey bars indicate the 5 min Arg stimulation.

All amino acids and pyruvate were included in at least three separate perfusion experiments. To combine the average effect of each amino acid on glucagon secretion from the perfused mouse pancreas, the fold change from baseline was calculated by dividing the average glucagon concentration during the amino acid stimulation with the preceding 5 min ‘wash period’ (this five min period is reffered to as baseline). Significance was assesed by comparing the average glucagon concentration during the amino acid stimulation with the average concentration during the preceding baseline using a paired t-test. Alanine (P=0.03), arginine (P<0.0001), and proline (P=0.03) increased glucagon secretion significantly from baseline. When evaluating the fold changes from baseline, the amino acids that increased glucagon secretion included the following: cysteine (8.4±3.8, P=0.08), alanine (2.1±0.4, P=0.03), glycine (1.7±0.3, P=0.2), proline (1.7±0.3, P=0.03), arginine (1.5±0.3, P<0.0001), and lysine (1.3±0.3, P=0.2) (Fig. 2) (Table 1). To our surprise, glutamine (1.0±0.05, P=0.2) did not stimulate glucagon secretion as previously described (9). The baseline values did not differ between the 19 amino acids (all comparisons P>0.9).

**Fig. 2.**
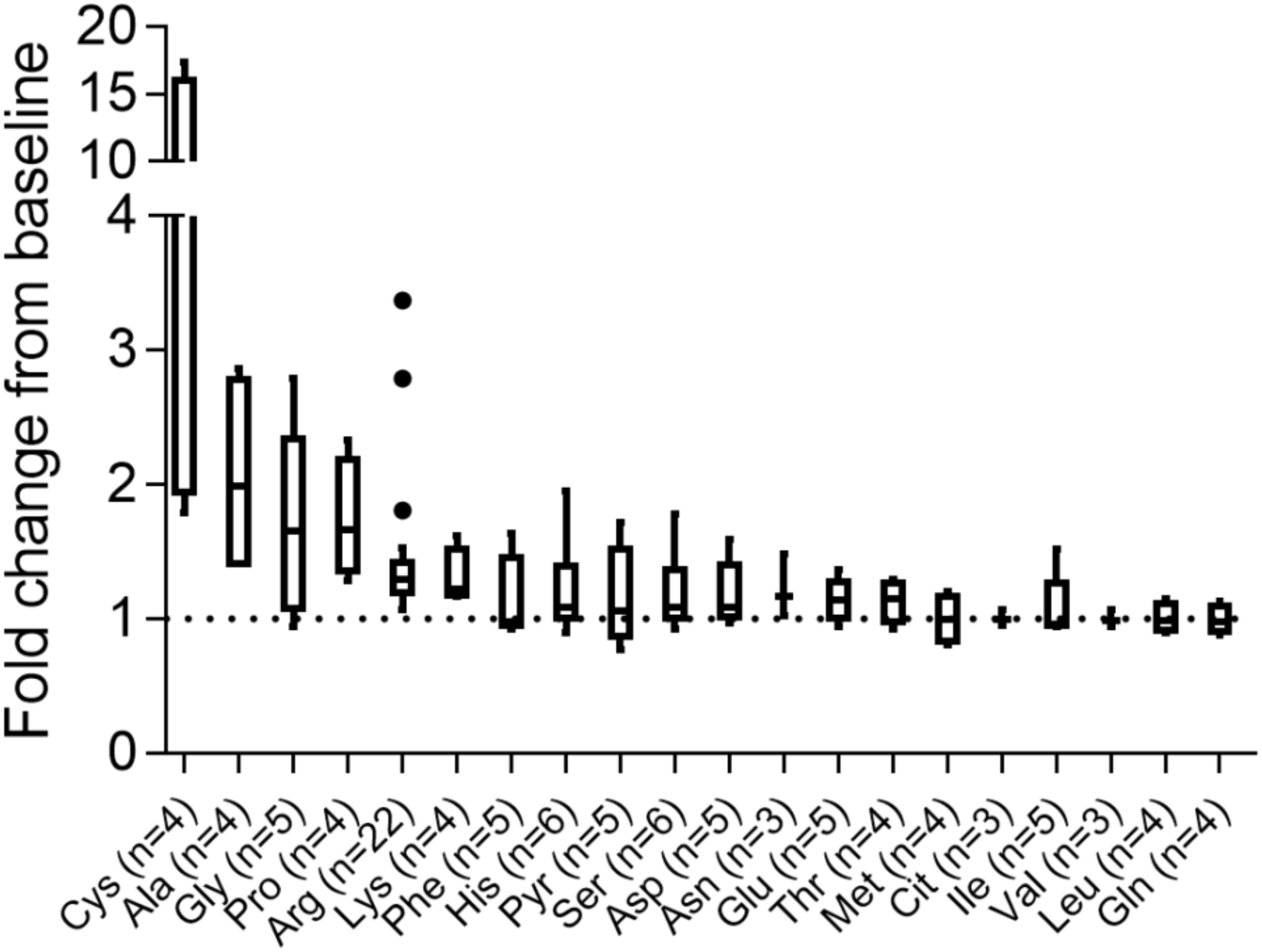
Amino acid induced glucagon secretion from the perfused mouse pancreas. Glucagon secretion is shown as fold change from baseline (mean±SEM), in effluent from the perfused pancreas in female C57BL/6JRj mice in response to; cysteine (Cys), alanine (Ala), glycine (Gly), proline (Pro), arginine (Arg), lysine (Lys), phenylananine (Phe), histidine (His), pyruvate (Pyr), serine (Ser), asparatate (Asp), asparagine (Asn), glutamate (Glu), threonine (Thr), methionine (Met), citrulline (Cit), isoleucine (Ile), valine (Val), leucine (Leu), and glutamine (Gln). The amino acids were administered by intra-arterial infusion at a concentration of 1 mM and the perfusate glucose concentration was kept at 3.5 mM.

To investigate whether the effect of these amino acids on glucagon secretion would be seen *in vivo* and to assess if the metabolism of those amino acids capable of stimulating glucagon secretion would be affected by glucagon receptor signaling, we divided the amino acids into four groups (mixture 1-4), according to their ability to stimulate glucagon secretion (table 2), and administered each of the four mixtures intraperitoneally three hours after glucagon receptor antagonist (GRA; 25-2648) or vehicle treatment in conscious mice.

### Glucagon secretion is stimulated equally by four isomolar mixtures of amino acids but differentially upon glucagon receptor antagonism

As expected, GRA treated mice showed increased plasma glucagon concentrations compared to vehicle treated mice (20.4±1.4 pmol/L vs. 6.9±0.4 pmol/L, P<0.0001). The basal glucagon concentrations did not differ between GRA treated mice across the four groups (mixture 1-4) (P>0.4), neither did the basal glucagon concentrations differ between the four vehicle groups (P>0.9).

Alanine, arginine, cysteine, and proline (mixture 1) increased plasma glucagon concentrations (shown as incremental area under the curve (_i_AUCs)) more in GRA than in vehicle treated mice (531.9±28.1 pmol/L × min vs. 225.8±25.0 pmol/L × min, P<0.0001) (Fig. 3*A* and *B*). Asparatate, glutamate, histidine, and lysine (mixture 2) also tended to increase glucagon concentrations more in GRA than in vehicle treated mice, however the difference was not significant (499.9±96.7 pmol/L × min vs. 251.4±24.8 pmol/L × min, P=0.06) (Fig. 3*C* and *D*). The increase in glucagon concentrations was significantly higher in GRA treated mice compared to vehicle treated mice upon administration of citrulline, methionine, serine, and threonine (mixture 3) (388.8±50.5 pmol/L × min vs. 200.6±33.8 pmol/L × min, P=0.01) (Fig. 3*E* and *F*), and after glutamine, isoleucine, leucine, and valine (mixture 4) (532.4±81.3 pmol/L × min vs. 225.9±42.0 pmol/L × min, P=0.007) (Fig. 3*G* and *H*).

**Fig. 3.**
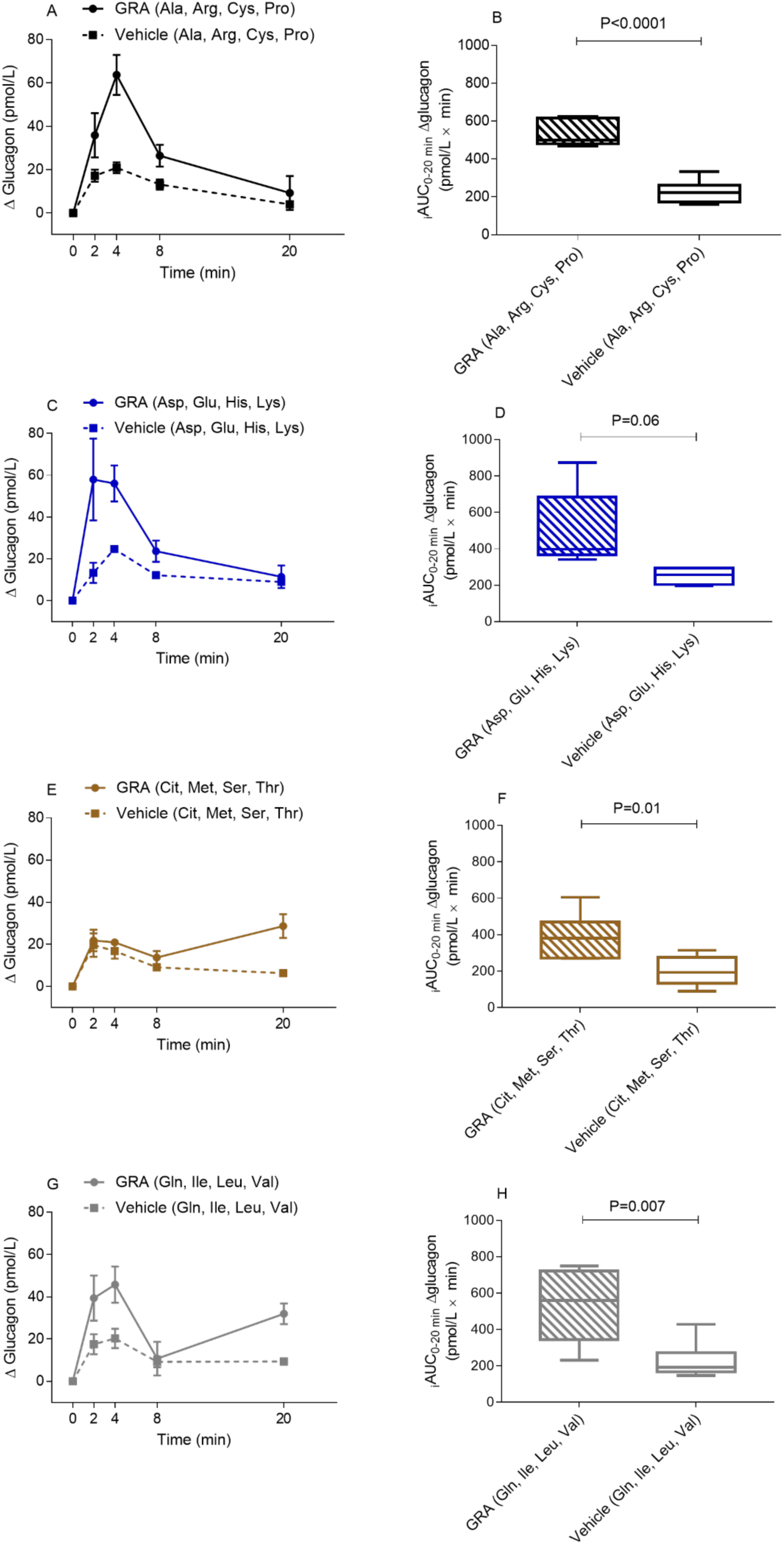
Glucagon secretion is stimulated equally by four isomolar mixtures of amino acids but differentially upon glucagon receptor antagonism. Plasma glucagon concentrations, shown as change from baseline in female C57BL/6JRj mice fasted for three hours and treated with vehicle (dotted lines and squares) or a glucagon receptor antagonist (GRA; 25-2648, 100 mg/kg body weight, administered by oral gavage) (solid lines and circles), upon intraperitoneal administration of a 7 µmol/g body weight amino acid mixture containing: (*A*) alanine (Ala), arginine (Arg), cysteine (Cys), and proline (Pro); (*C*) asparatate (Asp), glutamate (Glu), histidine (His), and lysine (Lys); (*E*) citrulline (Cit), methionine (Met), serine (Ser), and threonine (Thr); or (*G*) glutamine (Gln), isoleucine (Ile), leucine (Leu), and valine (Val). The basal glucagon concentrations did not differ between the four GRA groups (P>0.4), neither did the basal glucagon concentrations differ between the four vehicle groups (P>0.9). The incremental area under the curves (_i_AUC_0-20_ min) of plasma glucagon concentrations are shown as Tukey box-plots for the eight groups (*B, D, F*, and *H*). Data are presented as mean±SEM, n=4-6, P-value by unpaired t-test.

In contrast to the glucagon responses observed in our pancreas perfusion experiments, the glucagon concentrations (_i_AUCs) in response to the four amino acid mixtures *did not* differ when vehicle treated mice were compared (all comparisons P=0.9), neither did the glucagon responses differ when the _i_AUCs of GRA treated mice were compared (all comparisons P>0.5).

### Glucagon receptor antagonism differentially influences the disappearance of the individual amino acids

Three hours after GRA and vehicle treatment, plasma amino acid concentrations were increased in GRA treated mice compared to vehicle (4.4±0.2 mmol/L vs. 3.4±0.2 mmol/L, P=0.0001). Basal plasma amino acid concentrations were significant lower in GRA treated mice administered mixture 1 (3.3±0.2 mmol/L) when compared to GRA treated mice administered mixture 2 (4.9±0.2 mmol/L, P<0.0001), 3 (4.4±0.2 mmol/L, P<0.003), and 4 (5.3±0.2 mmol/L, P<0.0001). Basal amino acid concentrations were also lower in GRA treated mice administered mixture 3 when compared to GRA treated mice administered mixture 4 (P=0.02). No difference were observed between the remaining GRA groups (P>0.1). Basal amino acid concentrations were significant lower in vehicle treated mice administered mixture 1 compared to mixture 3 (2.8±0.2 mmol/L vs. 4.1±0.5 mmol/L, P=0.03). No difference were observed when the basal amino acid concentrations of the remaining vehicle groups were compared (P>0.1).

GRA treatment impaired the dissapearance of alanine, arginine, cysteine, and proline (mixture 1), as reflected by a significant greater _i_AUC in GRA treated mice compared to vehicle treated mice (33.4±3.1 mmol/L × min vs. 21.6±1.3 mmol/L × min, P=0.004) (Fig. 4*A* and *B*). The disappearance of citrulline, methionine, serine, and threonine (mixture 3) was also impaired in GRA treated mice when compared to vehicle (183.3±12.5 mmol/L × min vs. 138.2±13.5 mmol/L × min, P=0.04) (Fig. 4*E* and *F*). The disappearance of asparatate, glutamate, histidine, and lysine (mixture 2) did not differ between GRA and vehicle treated mice (31.0±5.1 mmol/L × min vs. 27.3±4.1 mmol/L × min, P=0.6) (Fig. 4*C* and *D*), and neither did the disappearance of glutamine, isoleucine, leucine, and valine (mixture 4) differ between GRA and vehicle treated mice (161.3±28.0 mmol/L × min vs. 190.3±128.0 mmol/L × min, P=0.4) (Fig. 4*G* and *H*).

**Fig. 4.**
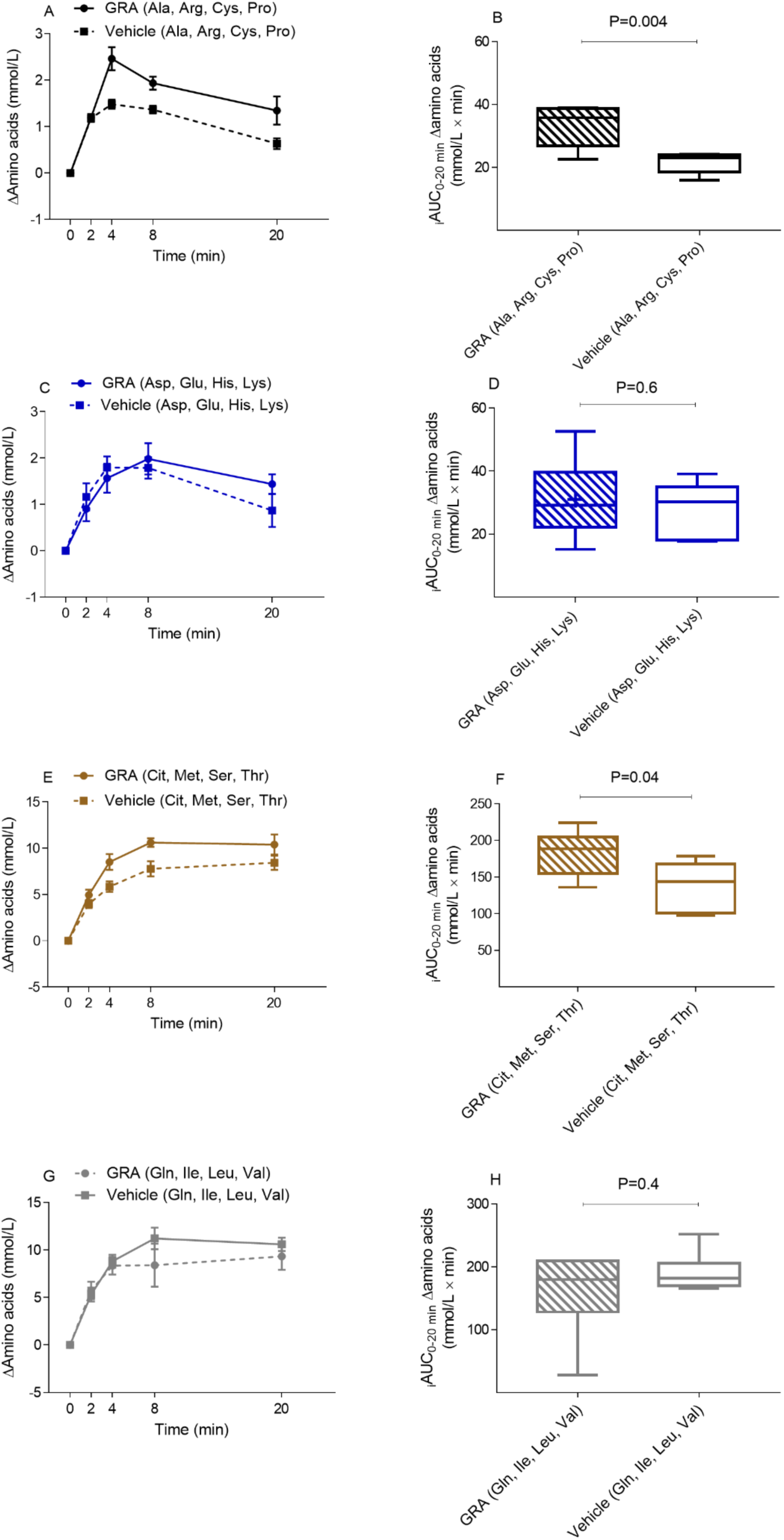
Glucagon receptor antagonism differentially influences the disappearance of the individual amino acids. Plasma amino acid concentrations, shown as change from baseline in female C57BL/6JRj mice fasted for three hours and treated with vehicle (dotted lines and squares) or a glucagon receptor antagonist (GRA; 25-2648, 100 mg/kg body weight, administered by oral gavage) (solid lines and circles) upon intraperitoneal administration of a 7 µmol/g body weight amino acid mixture containing: (*A*) alanine (Ala), arginine (Arg), cysteine (Cys), and proline (Pro); (*C*) asparatate (Asp), glutamate (Glu), histidine (His), and lysine (Lys); (*E*) citrulline (Cit), methionine (Met), serine (Ser), and threonine (Thr); or (*G*) glutamine (Gln), isoleucine (Ile), leucine (Leu), and valine (Val). The incremental area under the curves (_i_AUC_0-20_ min) of plasma amino acid concentrations are shown as Tukey box-plots for the eight groups (*B, D, F*, and *H*). Data are presented as mean±SEM, n=5-6, P-value by unpaired t-test.

### Administration of alanine, arginine, cysteine, and proline increases blood glucose concentrations in a glucagon receptor dependent manner

Before GRA and vehicle treatment blood glucose concentrations did not differ between GRA and vehicle treated mice (8.6±0.3 mmol/L vs. 8.9±0.2 mmol/L, P=0.4). GRA treatment resulted in decreased blood glucose concentrations (8.6±0.3 mmol/L vs. 6.6±0.2 mmol/L, P<0.0001), while blood glucose concentrations remained unchanged in vehicle treated mice (8.9±0.2 mmol/L vs. 9.1±0.1 mmol/L, P=0.2). The basal blood glucose concentrations did not differ between the four GRA groups (P>0.9), neither did the basal blood glucose concentrations differ between the four vehicle groups (P>0.9).

Administration of alanine, arginine, cysteine, and proline (mixture 1) increased blood glucose concentrations more in vehicle than in GRA treated mice, as reflected by a significant larger _i_AUC (42.8±5.8 mmol/L × min vs. 22.6±5.4 mmol/L × min, P=0.03) (Fig. 5*A* and *B*). Asparatate, glutamate, histidine, and lysine (mixture 2) increased blood glucose concentrations to a similar degree in GRA and vehicle treated mice (28.7±7.8 mmol/L × min vs. 30.2±7.8 mmol/L × min, P=0.9) (Fig. 5*C* and *D*). Blood glucose concentrations were also similar in GRA and vehicle treated mice upon administration of citrulline, methionine, serine, and threonine (mixture 3) (23.0±4.5 mmol/L × min vs. 34.7±7.8 mmol/L × min, P=0.2) (Fig. 5*E* and *F*), and glutamine, isoleucine, leucine, and valine (mixture 4) (32.8±7.3 mmol/L × min vs. 19.7±2.8 mmol/L × min, P=0.1) (Fig. 5*G* and *H*).

**Fig. 5.**
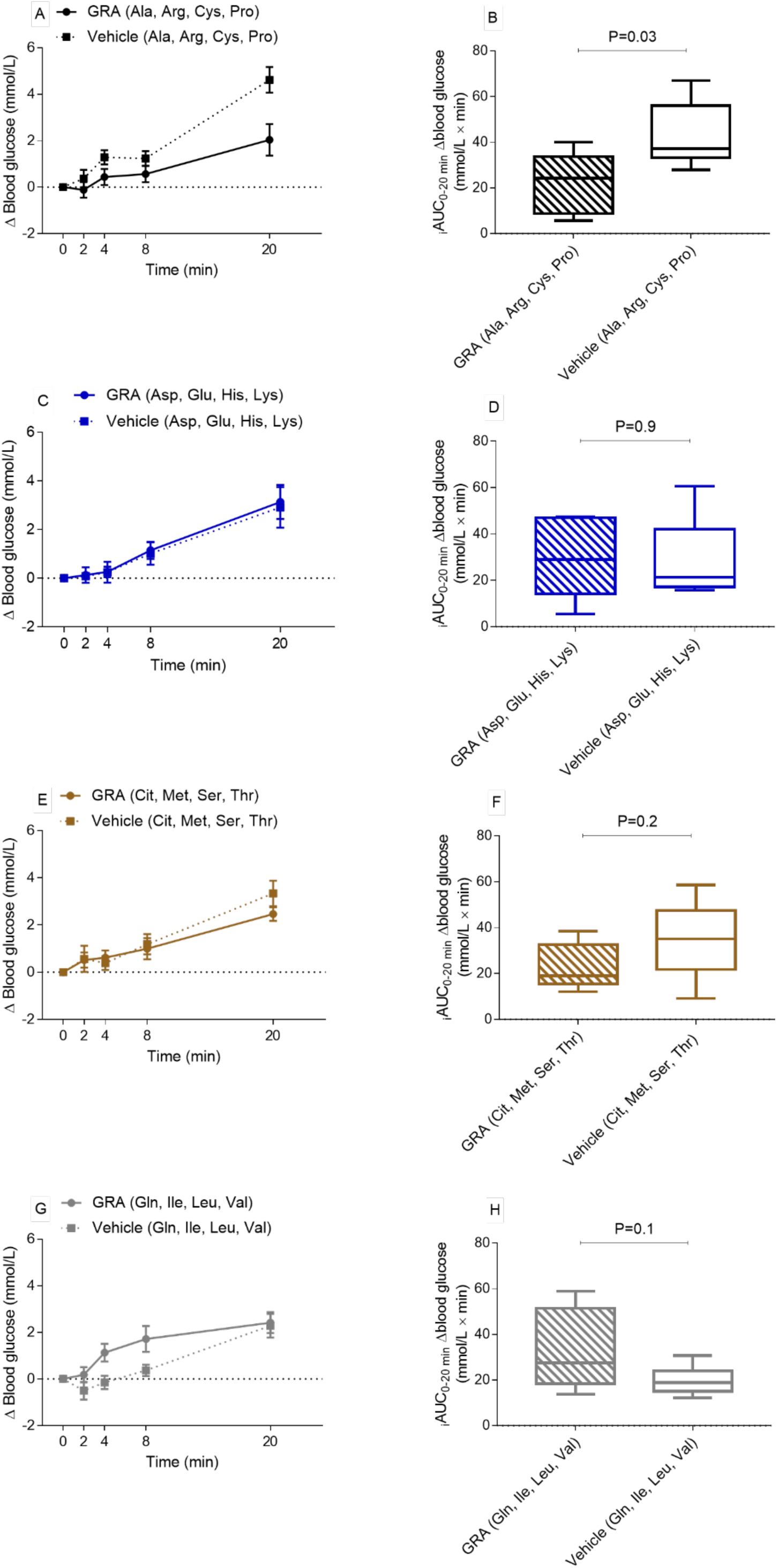
Administration of alanine, arginine, cysteine, and proline increases blood glucose concentrations in a glucagon receptor dependent manner. Blood glucose concentrations, shown as change from baseline in female C57BL/6JRj mice fasted for three hours and treated with vehicle (dotted lines and squares) or a glucagon receptor antagonist (GRA; 25-2648, 100 mg/kg body weight, administered by oral gavage) (solid lines and circles), upon intraperitoneal administration of a 7 µmol/g body weight amino acid mixture containing: (*A*) alanine (Ala), arginine (Arg), cysteine (Cys), and proline (Pro); (*C*) asparatate (Asp), glutamate (Glu), histidine (His), and lysine (Lys); (*E*) citrulline (Cit), methionine (Met), serine (Ser), and threonine (Thr); or (*G*) glutamine (Gln), isoleucine (Ile), leucine (Leu), and valine (Val). The basal blood glucose concentrations did not differ between the four GRA groups (P>0.9), neither did the basal blood glucose concentrations differ between the four vehicle groups (P>0.9).The incremental area under the curves (_i_AUC_0-20_ min) of blood glucose concentrations are shown as Tukey box-plots for the eight groups (*B, D, F*, and *H*). Data are presented as mean±SEM, n=5-6, P-value by unpaired t-test.

### Glucagon receptor antagonism enhances the insulin secretion stimulated by branched chained amino acids and glutamine

Plasma insulin concentrations did not differ between GRA and vehicle treated mice in the basal state (0.6±0.1 ng/mL vs. 0.8±0.1 ng/mL, P=0.2). The basal insulin concentrations did not differ between the four GRA groups (P>0.4), neither did the basal insulin concentrations differ between the four vehicle groups (P>0.9).

Plasma insulin concentrations (_i_AUCs) were similar in GRA and vehicle treated mice upon administration of mixture 1, 2, and 3; alanine, arginine, cysteine, and proline (mixture 1) (5.8±2.7 ng/mL × min vs. 2.3±1.4 ng/mL × min, P=0.3) (Fig. 6*A* and *B*); asparatate, glutamate, histidine, and lysine (mixture 2) (2.5±1.5 ng/mL × min vs. 1.4±1.4 ng/mL × min, P=0.6) (Fig. 6*C* and *D*); citrulline, methionine, serine, and threonine (mixture 3) (4.7±1.9 ng/mL × min vs. 3.5±1.6 ng/mL × min, P=0.6) (Fig. 6*E* and *F*). In contrast, insulin concentrations were significantly higher in GRA treated mice upon administration of glutamine, isoleucine, leucine, and valine (mixture 4) (12.6±2.7 ng/mL × min vs. 4.4±1.8 mg/mL × min, P=0.03) (Fig. 6*G* and *H*).

**Fig. 6.**
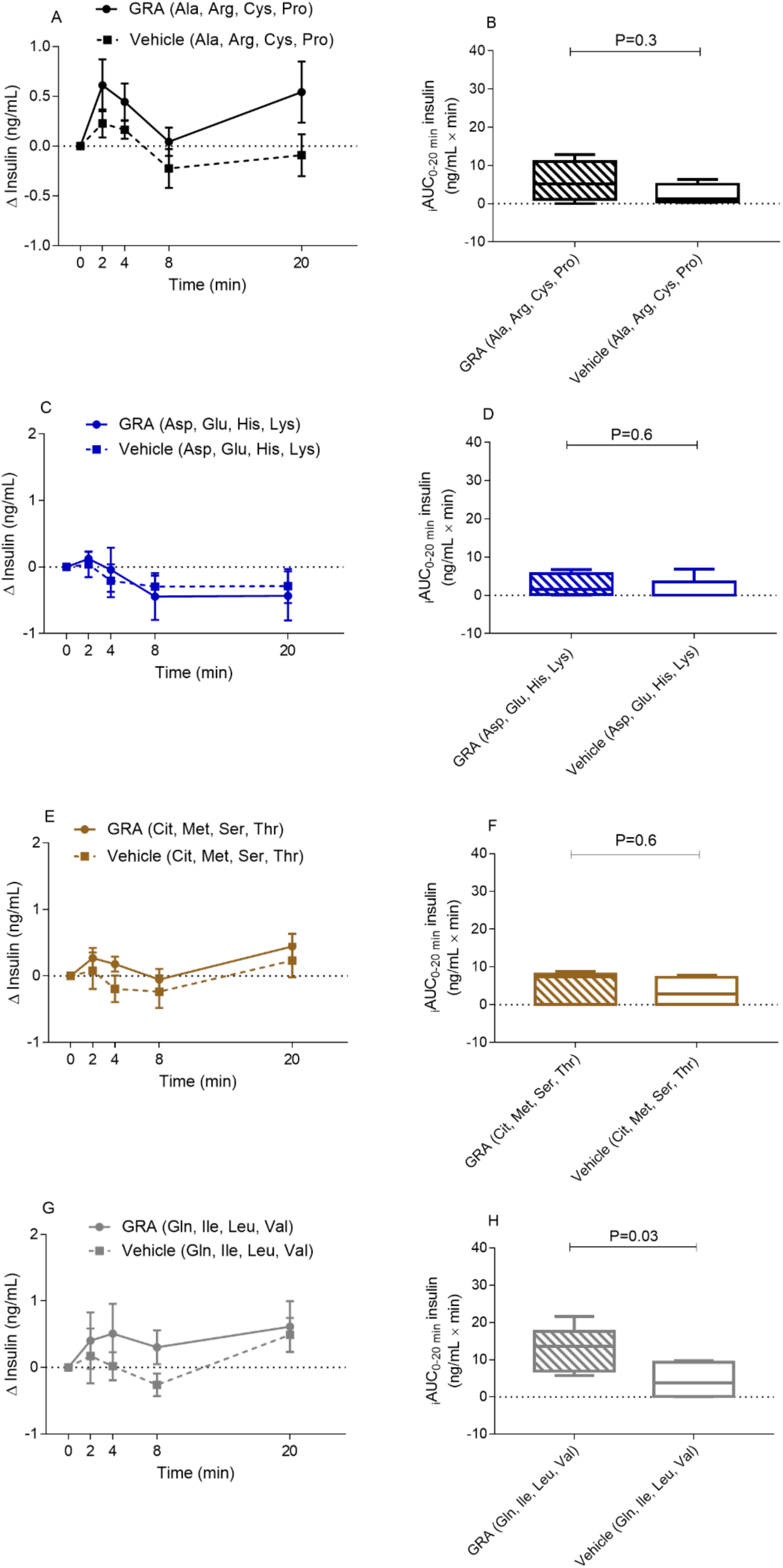
Glucagon receptor antagonism enhances the insulin secretion stimulated by branched chained amino acids and glutamine. Plasma insulin concentrations, shown as change from baseline in female C57BL/6JRj mice fasted for three hours and treated with vehicle (dotted lines and squares) or a glucagon receptor antagonist (GRA; 25-2648, 100 mg/kg body weight, administered by oral gavage) (solid lines and circles), upon intraperitoneal administration of a 7 µmol/g body weight amino acid mixture containing: (*A*) alanine (Ala), arginine (Arg), cysteine (Cys), and proline (Pro); (*C*) asparatate (Asp), glutamate (Glu), histidine (His), and lysine (Lys); (*E*) citrulline (Cit), methionine (Met), serine (Ser), and threonine (Thr); or (*G*) glutamine (Gln), isoleucine (Ile), leucine (Leu), and valine (Val). The basal insulin concentrations did not differ between the four GRA groups (P>0.4), neither did the basal insulin concentrations differ between the four vehicle groups (P>0.9). The incremental area under the curves (_i_AUC_0-20_ min) of plasma insulin concentrations are shown as Tukey box-plots for the eight groups (*B, D, F*, and *H*). Data are presented as mean±SEM, n=4-6, P-value by unpaired t-test.

No difference in insulin concentrations was observed in response to the four amino acid mixtures when the _i_AUCs of vehicle treated mice were compared (all comparisons P>0.7). When the _i_AUCs of GRA treated mice were compared the insulin concentrations were significantly higher after mixture 4 compared to mixture 2 (P=0.049), there was no difference between the remaining groups (all comparisons P>0.1).

## Discussion

Here we demonstrate that alanine, arginine, and proline stimulate glucagon secretion in the perfused mouse pancreas. Cysteine, glycine, and lysine did not reach statistical significance but resulted in numerically higher glucagon responses compared to baseline. The remaining amino acids including the BCAAs and glutamine had no effect on glucagon secretion in female mouse pancreata at the tested dose. The ability of amino acids to stimulate glucagon secretion differs, and the amino acid induced glucagon secretion differs between species and experimental conditions. In isolated mouse islets prolonged administration of amino acids lead to alpha cell proliferation (57), and alanine, glutamate, glutamine, and leucine have in particular been associated with alpha cell proliferation (15, 30). In perifused mouse islet, alanine (10 mM) has been shown to be a more potent stimulant of glucagon secretion than glutamine (10 mM) (10), however in another perifused islet study by the same group, glutamine (10 mM) and arginine (1 mM) were both found to be potent stimulators of glucagon secretion (9). In several species alanine, arginine, proline, glycine, and lysine seem to be potent stimulators of glucagon secretion, whereas the BCAAs have not been reported to influence secretion (17, 33, 52), except for one study that found 20 min of valine and leucine infusion to stimulate glucagon secretion in the perfused rat pancreas (3) (table 3). We administered 1 mM of each amino acid, which is substantially higher than the circulating concentration of any amino acid (which ranges from 6 µmol/L to 600 µmol/L in C57BL/6JRj mice (20)) (table 1), but lower than the 10-25 mM applied in other studies done in isolated islets, in which the majority of amino acids have been found to stimulate glucagon secretion (3, 9, 10, 17).

**Table 3.** Amino acid induced glucagon secretion in four studies evaluating glucagon secretion in response to individual amino acids. - indicates that the amino acid did not increase glucagon secretion, X indicates that the amino acid did result in an increase in glucagon secretion, and blank indicates that the amino acid was not administered.

| Amino acid | Stimulation of the perfused rat pancreas with individual amino acid (25 mM) (glucose 4.4 mM) (3). | Stimulation of perfused islet from high protein fed rats with individual amino acid (20 mM) (glucose 1.7 mM) (17). | Intravenous infusion of individual amino acid (3 mmol/kg) in sheep (33). | Intravenous infusion of individual amino acids (1 mmol/kg) in dogs (52). |
| --- | --- | --- | --- | --- |
| Cysteine | - | - |  | X |
| Alanine | X | - | X | X |
| Glycine | X | - | X | X |
| Proline |  | X | X | X |
| Arginine | X | X | X | X |
| Lysine |  | X | X | X |
| Phenylalanine | - | - | X | X |
| Histidine |  | - | X | X |
| Pyruvate |  |  |  | - |
| Serine | X | - | X | X |
| Asparatate |  | - | - | X |
| Asparagine | X | - | X | X |
| Glutamate | X | - | - | X |
| Threonine |  | - | X | X |
| Methionine |  | - | X | X |
| Citrulline | - | - |  |  |
| Isoleucine |  | - | - | - |
| Valine | X | - | - | - |
| Leucine | X | - | - | - |
| Glutamine |  | X | X | X |

The glucagon responses observed in the perfused pancreas were not reflected in our *in vivo* studies, as we found all of the amino acid mixtures to stimulate glucagon secretion equally, although the dynamics of glucagon secretion differed between the four mixtures in GRA treated mice. GRA enhanced the glucagon secretion 2 and 4 min after administration of amino acid mixture 1 and 2, whereas GRA enhanced only glucagon secretion at 20 min after mixture 3. After mixture 4, GRA enhanced glucagon secretion both 2, 4 and 20 min after administration. This differential glucagon secretion may be due to intracellular processing of certain amino acids leading to activation of pathways that induce glucagon secretion. The ability of the four amino acid mixtures to stimulate glucagon secretion equally in vehicle treated mice may be due to rapid conversion of non-glucagonotrophic amino acids to glucagonotrophic amino acids, or *vice versa*, in the circulation. Additionally, by administering 7 µmol/g body weight of the amino acid mixtures we increased plasma concentrations of the administered amino acids to supraphysiological levels, and may thus have stimulated glucagon secretion to a point where a potential difference among the amino acids disappeared, and some amino acids having little, if any, effect on glucagon secretion alone may have enhanced the glucagon secretion stimulated by other amino acids (50). GRA potentiated the glucagon secretion in response to all four mixtures. This effect of GRA raises the question of autocrine glucagon regulation, and since glucagon receptor mRNA transcripts has been found in alpha cells it is possible that glucagon binds its own receptor and regulates alpha cell secretion (37, 41). Within the islets, glucagon concentrations may be sufficient to activate glucagon-like peptide-1 (GLP-1) receptors present on beta cells, but when testing our GRA *in vitro*, we found that GRA inhibited glucagon action whether on the glucagon or the GLP-1 receptors (21). GRA increased plasma amino acid concentrations, and we believe that the GRA augmented glucagon secretion may be due to feedback mechanisms involving amino acids and/or beta- and delta-cell secretory products. The augmented insulin secretion in GRA treated mice in response to mixture 4 is surprising since glucagon has been shown to induce insulin secretion (9, 59, 70). We therefore expected GRA treatment to, if anything, decrease insulin secretion. Administration of mixture 4, surprisingly, tended to result in higher blood glucose concentrations in GRA treated mice compared to vehicle treated mice, which might explain the increased insulin concentrations. Upon administration of mixture 3 and 4 (and 1 in GRA treated mice) there was a tendency for insulin concentrations to increase at 20 min, this may reflect intracellular processing of amino acids leading to activation of pathways that induce insulin secretion. Intracellular amino acid metabolism might also explain the increase in glucagon concentrations observed in GRA treated mice at 20 min upon mixture 3 and 4. Additionally, some of the administered amino acid could have been converted to other amino acids causing the second peak in insulin and glucagon concentrations. To confirm and elucidate the conversion of amino acids in the circulation, one would have to measure plasma concentration of the individual amino acids at various time-points following administration. This was however not technical possible due to volume restriction (ethical considerations) and methodology of measurement techniques.

During conditions of hyperglucagonemia, plasma amino acid concentrations decrease and alanine, arginine, proline, glycine, lysine, and threonine seems to exhibit the largest decreases (7, 18, 23, 39, 42, 44, 51, 53). The opposite is observed during conditions of glucagon action deficiency, where plasma concentrations of the same amino acids (alanine, arginine, proline, glycine, lysine, and threonine) increase while the concentrations of BCAAs and aromatic amino acids (phenylalanine, tryptophan, and tyrosine) seem to increase only during conditions of chronic glucagon deficiency (6, 7, 20, 30, 57, 69). Glucagon may acutely affect the metabolism of these amino acids without changing the plasma concentration, as observed for phenylalanine (60). It is important to note that during both chronic glucagon excess (glucagonoma patients (2, 4)) and deficiency (subjects with glucagon receptor inactivation mutations (38)) all amino acids eventually become affected.

Glucagon stimulates endogenous protein catabolism (27, 32, 54, 68) and when acting on its main target, i.e. the liver, glucagon may lead to a net hepatic output of BCAAs (42) which are not catabolized by the liver (8, 14). It is therefore important to keep in mind that we measured total plasma amino acid concentrations and thus might have underestimated a decrease in certain amino acids while missing an increase in other amino acids. We found that the disappearance of alanine, arginine, cysteine, and proline (mixture 1) and of citrulline, methionine, serine, and threonine (mixture 3) was impaired upon pharmacological disruption of glucagon receptor signaling (GRA; 25-2648, 100 mg/kg body weight), while the disappearance of asparatate, glutamate, histidine, and lysine (mixture 2) and the BCAAs and glutamine (mixture 4) was unaffected. Glucagon regulates the expression of hepatic amino acid transporters (13, 16, 30, 36, 55, 57, 61, 69), and GRA treatment (for a period >1 hour) may downregulate specific amino acid transporters and hence impair the disappearance of the amino acids transported by glucagon regulated amino acid transporters. In line with the decreased disappearance of mixture 1 (presumably due to decreased hepatic amino acid uptake) we observed that GRA diminished the increase in blood glucose concentrations upon administration of mixture 1, the same tendency was observed upon administration of mixture 3, which disappearance was also impaired in GRA treated mice. The decreased amino acid uptake may result in decreased hepatic availability of substrates for glucose production. Glucagon is a known regulator of gluconeogenesis (29), and disruption of glucagon signaling results in downregulation of rate-limiting enzymes of gluconeogenesis such as phosphoenolpyruvate carboxykinase (69). Alanine, arginine, and proline are gluconeogenic amino acids (33, 46, 67), and the diminished increase in blood glucose concentrations after GRA treatment may reflect decreased gluconeogenic activity. In line with this, alanine failed to increase blood glucose concentrations in proglucagon knockout mice, suggesting that glucagon might be needed for alanine-induced glucose production (10).

A clear limitation to our study is the lack of individual amino acid administration *in vivo*, and this obviously limits the interpretations of our *in vivo* results. During physiological conditions amino acids are never present without other amino acids, however our study, being a mechanistic study aiming to elucidate the role of individual amino acids in the liver-alpha cell axis, clearly depend on administration of individual amino acids. Another limitation to the study is the absolute differences in plasma amino acid concentrations in response to the isomolar amino acid mixtures. This may be related to measurement technique (recovery) or conversion rates *in vivo*. Finally, the glucagon receptor antagonist was administered three hours prior to administration of the amino acid mixtures, during which period a new set point may have been established, as indicated by increased plasma amino acid and glucagon concentrations and decreased blood glucose concentrations, which may have influenced our results.

Our data show that alanine, arginine, and proline are glucagonotropic in the perfused mouse pancreas, while pharmacological inhibition of glucagon receptor signaling impairs the disappearance of alanine, arginine, cysteine, and proline (mixture 1) and citrulline, methionine, serine, and threonine (mixture 3). Our results, when reviewed together with the existing literature regarding this subject, suggest that alanine, arginine, and proline are the main amino acids participating in the liver-alpha cell axis in mice.

## Acknowledgements

We thank Jesper Lau, Novo Nordisk A/S, Måløv, Denmark, for providing the glucagon receptor antagonist 26-2548.

## Author contributions

Conceived and designed research: K.D.G., N.J.W.A., and J.J.H. Performed experiments: K.D.G., S.L.J., and S.A.S.K. Analyzed data: K.D.G., S.L.J., and N.J.W.A. Interpreted results of experiments: K.D.G., S.L.J., N.J.W.A., and J.J.H. Prepared figures: K.D.G. and N.J.W.A. Drafted manuscript: K.D.G., and N.J.W.A. Edited and revised manuscript: S.L.J., S.A.S.K., J.P., and J.J.H. Approved final version of manuscript: K.D.G., S.L.J., S.A.S.K., J.P., N.J.W.A., and J.J.H.

---

**Supplementary table 1.** The amino acids used in the pancreas perfusion experiments and the *in vivo* experiments.

| Amino Acid | Catalog number | Company |
| --- | --- | --- |
| Alanine | A7469-25G | Sigma-Aldrich |
| Arginine | A8094-25G | Sigma-Aldrich |
| Asparagine | A8381-100G | Sigma-Aldrich |
| Asparatate (Aspartic acid) | 11189-100G | Sigma-Aldrich |
| Cysteine | C7880-100G | Sigma-Aldrich |
| Glutamate (Glutamic Acid) | G5667-100G | Sigma-Aldrich |
| Glutamine | G5792-100G | Sigma-Aldrich |
| Glycine | G5417-100G | Sigma-Aldrich |
| Histidine | H6034-25G | Sigma-Aldrich |
| Isoleucine | LI7403-25G | Sigma-Aldrich |
| Leucine | L8912-25G | Sigma-Aldrich |
| Lysine | L8662-25G | Sigma-Aldrich |
| Methionine | M5308-25G | Sigma-Aldrich |
| Phenylalanine | P5482-25G | Sigma-Aldrich |
| Proline | P5607-25G | Sigma-Aldrich |
| Pyruvate (Sodium Pyruvate) | P2256-25G | Sigma-Aldrich |
| Serine | S4311-25G | Sigma-Aldrich |
| Threonine | T8441-25G | Sigma-Aldrich |
| Valine | V0513-25G | Sigma-Aldrich |

**Supplementary figure 1.**
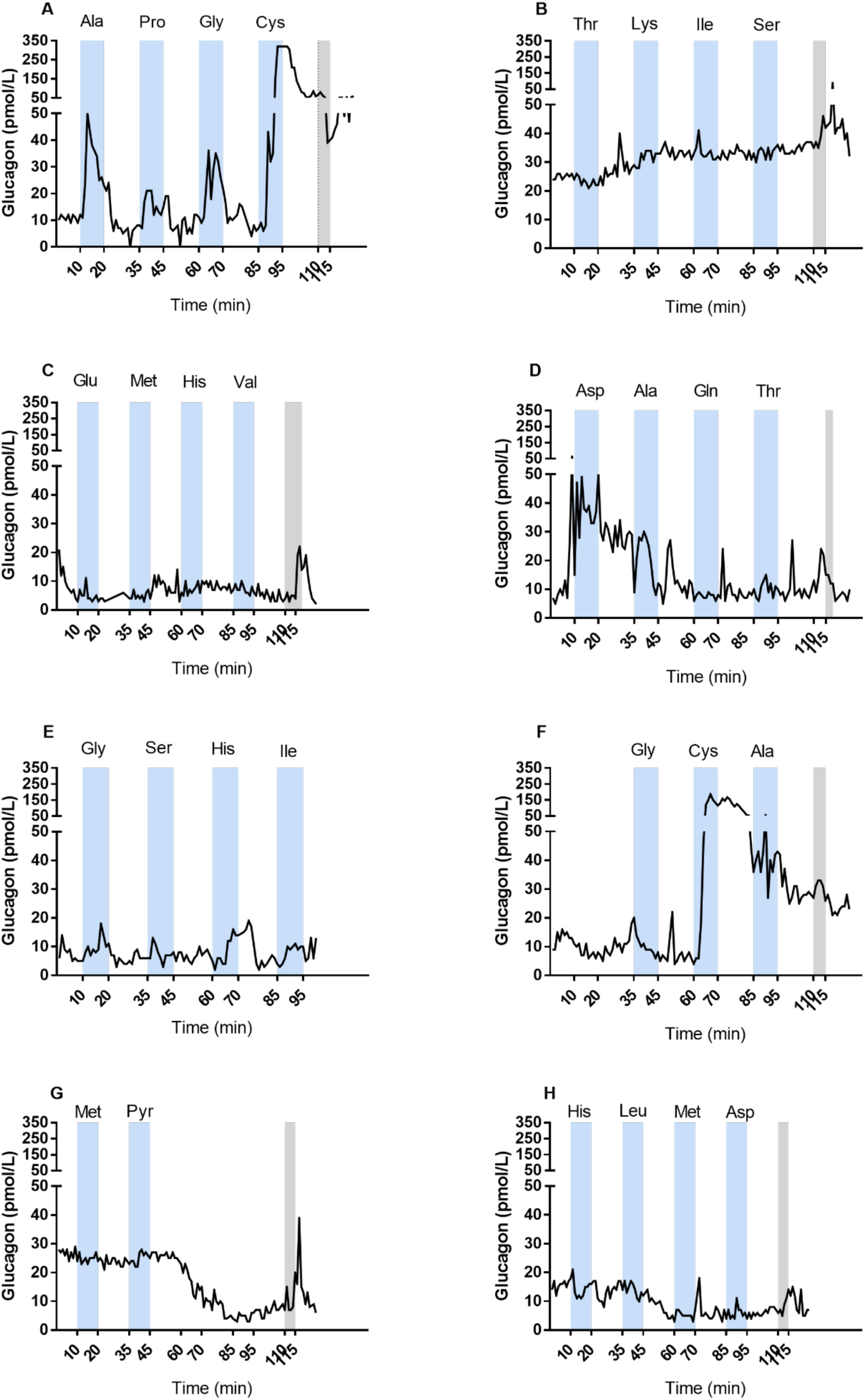
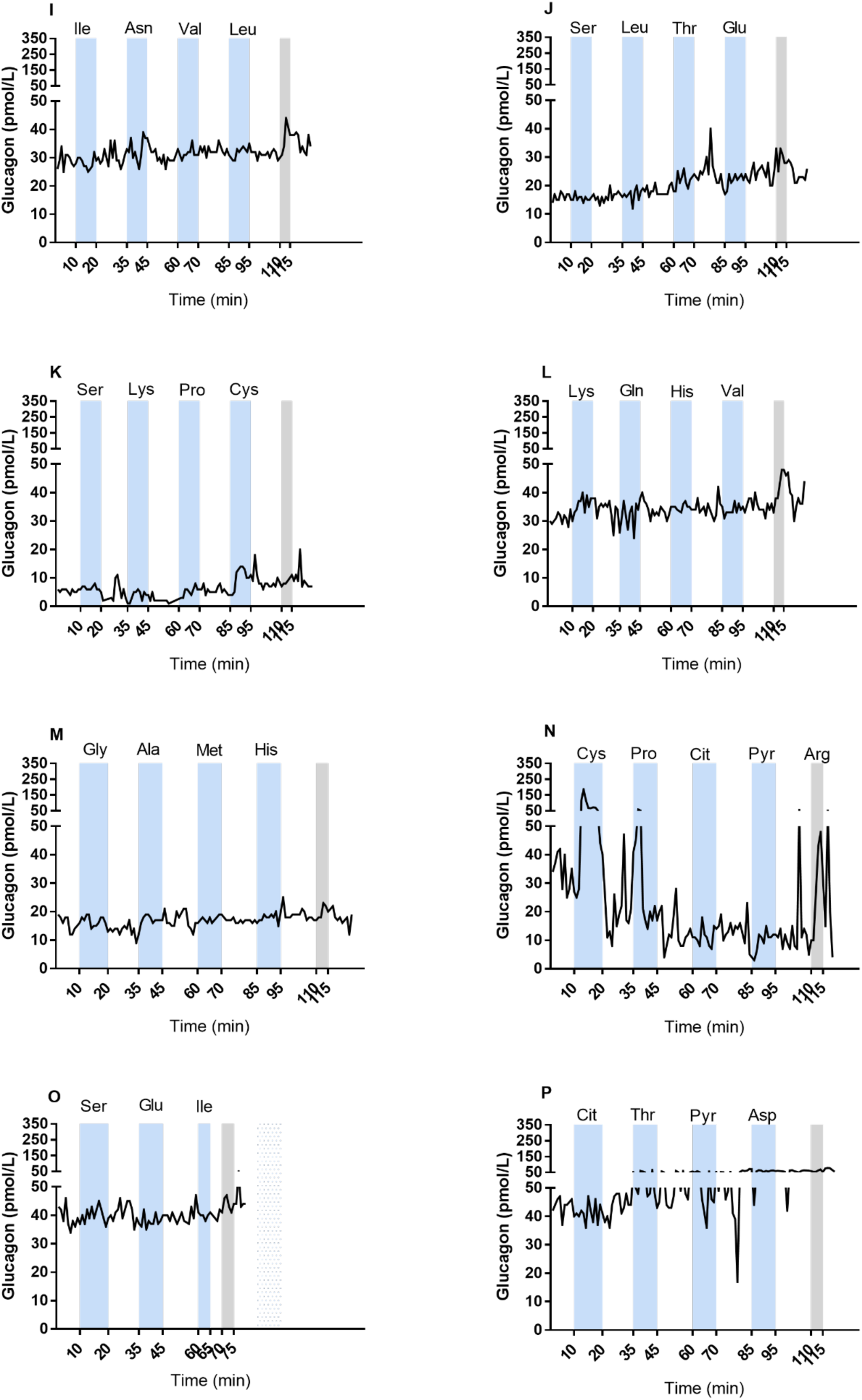
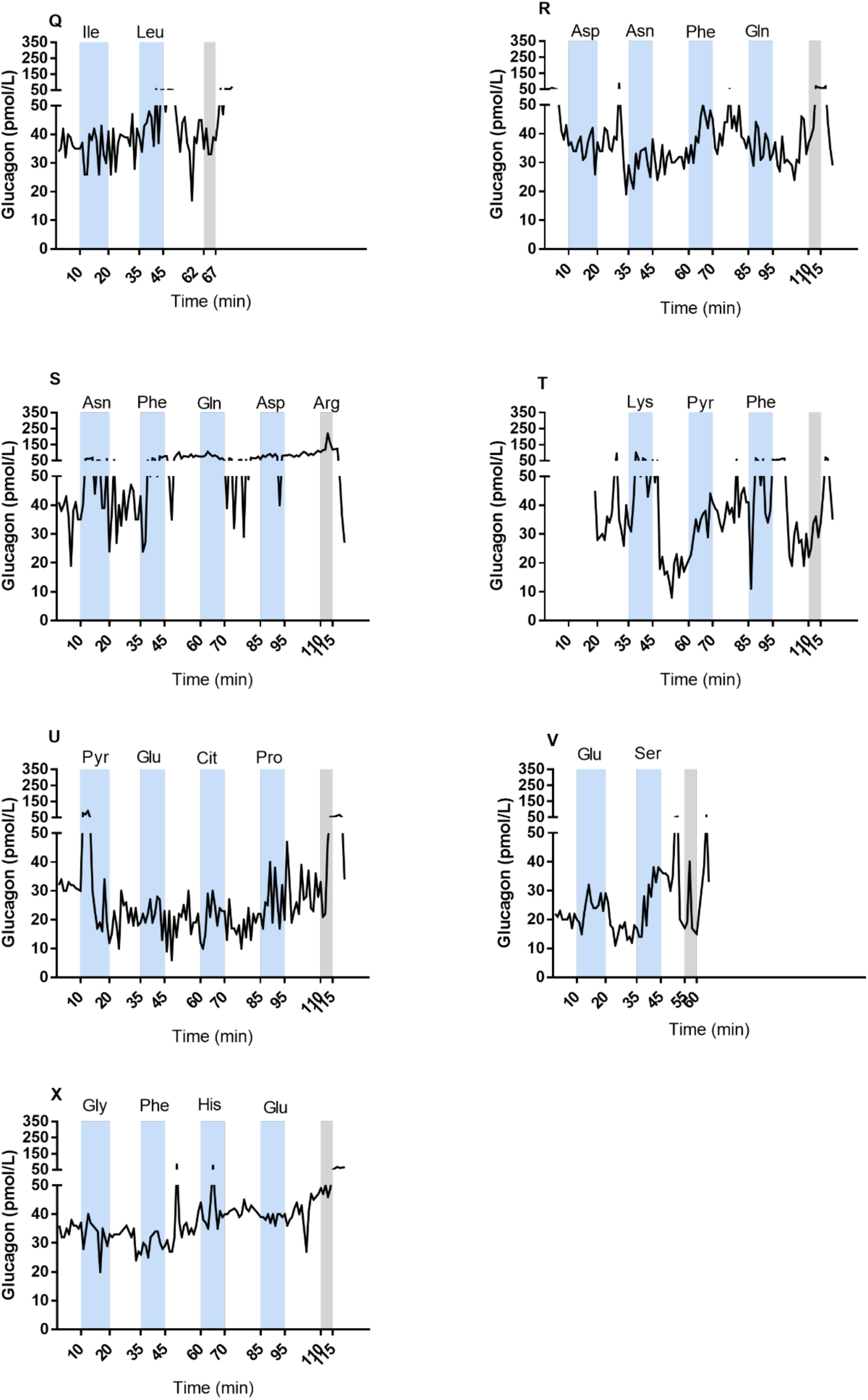
Glucagon concentrations in effluent from the perfused pancreas in non-fasted female C57BL/6JRj mice in response to individual amino acids; cysteine (Cys), alanine (Ala), glycine (Gly), proline (Pro), arginine (Arg), lysine (Lys), phenylananine (Phe), histidine (His), pyruvate (Pyr), serine (Ser), asparatate (Asp), asparagine (Asn), glutamate (Glu), threonine (Thr), methionine (Met), citrulline (Cit), isoleucine (Ile), valine (Val), leucine (Leu), and glutamine (Gln). All amino acids were administered by intra-arterial infusion at a concentration of 1 mM and the perfusate glucose concentration was kept at 3.5 mM. The blue bars indicate the amino acid stimulations and the grey bars indicate the Arg stimulation.

